# Basic region variants of the MAX b-HLH-LZ preferentially form heterodimers with the MYC b-HLH-LZ to bind the E-box rather than binding as homodimers

**DOI:** 10.64898/2026.04.01.715400

**Authors:** Vincent Roy, Martin Montagne, Pierre Lavigne

## Abstract

The MYC associated factor X (MAX) is the heterodimeric partner of the MYC paralogs (MYC, MYCN and MYCL). When deregulated, high level of the MYC paralogs contribute to all aspects of tumorigenesis and tumor growth. MAX can also heterodimerize with the MXD proteins, MNT and MGA. Heterodimerization and sequence specific DNA binding to the E-Box sequences at gene promoters is controlled by their heterodimerization with the MAX b-HLH-LZ. As a heterodimer with MAX, MYC proteins activate genes involved in cell metabolism, growth and proliferation whereas MXD proteins, MNT and MGA repress them. MAX can also bind to the E-Bos sequence as a homodimer. Being devoid of a transactivation domain it can act as an antagonist of the MYC/MAX heterodimers. Variants of MAX have been reported to be linked to cancer. These variants are either not expressed, inactivated or lead to missense mutations. This has led to the notion that MAX may have a tumor suppressor role. Here, we characterize three of those variants with missense mutations in the basic region, *i.e.* E32K, R35P and R35C. We analyzed their heterodimerization with the b-HLH-LZ of MYC and their DNA binding properties as homo-and heterodimers. The R35C variant b-HLH-LZ was found to have a markedly increased affinity for the b-HLH-LZ of MYC. We also observed that all three b-HLH-LZ variants have a lower affinity as homodimers for the E-Box than the WT. This was shown to lead to a preferential binding of all the heterodimeric b-LHLH-LZ to the E-Box. This effect is exacerbated in the case of the R35C variant. We argue that this preferential binding of MYC as heterodimers with these variants to E-Box sequences could contribute to tumorigenesis. Hence, our results suggest that, mechanistically, the MAX homodimer bound to the E-Box could act as a tumor suppressor.

## MATERIALS AND METHODS

### Molecular modeling

The open source version 1.7.6.0 of Pymol was used for modeling and molecular rendering [1]. The crystal structure of the MAX homodimer bound to the E-Box (1HLO [2]) was used as a template for the generation of the models. The variants were generated using the mutagenesis function in the wizard. The conformation of the K^32^ side chain was manually set in order to avoid introducing steric clashes with DNA.

### Protein expression and purification

The cDNA, coding for the MAX b-HLH-LZ (Max* hereafter, residues 22-103, UniProt entry P61244-1) to which are added the GSGC residues in c-terminal, inserted in the pET3a vector was already available in the laboratory [3] and was used as a template to generate the plasmids with inserts coding for each of the mutants (E32K, R35C and R35P) through quick-change PCR with Q5 DNA polymerase and DpnI from New England Biolabs. The primers used were purchased from IDT DNA, their sequences are listed in Table S1. Sequence for each construct was confirmed by Sanger sequencing at the Plateforme de séquençage SANGER – Centre de recherche du CHU de Québec – Université Laval. The primary structure for the basic region of each construct is given in **Fig. 2A**.

The MYC b-HLH-LZ (Myc*), the Max*^WT^ b-HLH-LZ and its variants were expressed and purified as previously described [3,4] After lyophilisation, the b-HLH-LZs were kept at -20°C and solubilised in Myc buffer (50 mM NaCl, 50 mM NaH_2_PO_4_ pH 5.5) for Myc* or PBS for Max* at a final concentration of 1 mM before use.

### Circular dichroism

All circular dichroism (CD) measurements were performed on a Jasco J-810 spectropolarimeter equipped with a Peltier-type thermostat. The instrument was routinely calibrated using an aqueous solution of *d*-10-(+)-camphorsulfonic acid at 290.5 nm. Samples were prepared as follows: Max* (either WT or a variant) was diluted in 100 µl 2X CD buffer (40 mM KCl, 11.4 mM K_2_HPO_4_, 28.6 mM KH_2_PO_4_, pH 6.8) and the volume adjusted to 106 µl with PBS. 10 µl TCEP 16 mM were added, and the volume further adjusted to 192 µl with ddH_2_O before samples were incubated overnight at room temperature. After reduction, Myc* was added and the volume adjusted to 198 µl with Myc buffer (Na_2_HPO_4_ 0.95 mM, NaH_2_PO_4_ 49.05 mM, 50 mM NaCl, pH 5.5).

The DNA complexes were prepared as follows. After a 10 minutes incubation of the protein samples at room temperature, 0, 1 or 2 µl of 2 mM of specific or non-specific DNA duplexes in 10 mM Tris pH 8.0 were added and the volume adjusted to 200 µl with 10 mM Tris pH 8.0. The strands of the specific probe were: 5’-ATT ACC <u>CAC GTG</u> TCC T*AC-3’ and 5’-GTA GGA <u>CAC GTG</u> GGT* AAT-3’ (with the E-box sequence underlined) and the non-specific probe: 5’-ATT ACC TCC GGA TCC T*AC-3’ and 5’-GTA GGA TCC GGA GGT* AAT-3’ (Integrated DNA Technologies). Samples were further incubated for 10 minutes at room temperature and transferred to a 1 mm path length quartz cuvette. All spectra were recorded from 250 to 195 nm at 0.1 nm intervals by accumulating 10 spectra at 25 °C. Thermal denaturations were recorded at 222 nm from 5 to 95 °C at a heating rate of 1 °C/min. CD signal for spectra and thermal denaturations was corrected by substracting the signal from corresponding spectra or thermal denaturation either for buffer alone or the appropriate DNA duplex. CD signal was then converted to mean residue ellipticity using the following formula [5]: [θ] = δ · MRW/(10 · *c* · *l*) where [θ] is the mean residue ellipticity in deg · cm^2^ · dmol^-1^, δ is the CD signal in millidegrees, MRW is the mean residue weight, *c* is the concentration in mg/ml and *l* is the pathlength in mm. For the heterodimers, the concentration used was the sum of Max* and Myc* and the MRW was determined using a weighted average.

## INTRODUCTION

The *max* gene (MYC-associated factor X) encodes a b-HLH-LZ transcription factor that promotes specific DNA binding of the MYC, MYCN and MYCL paralogs to the canonical E-box (CACGTG) as well as the activation of the transcription of genes controlling many cellular programs such as metabolism, growth and proliferation [6]. MAX can also heterodimerize with the MXD proteins (MXD1, MXD2, MXD3 and MXD4), MNT and MGA to antagonize the MYC/MAX heterodimer [7–11]. This results in the repression of the transcription of genes activated by MYC/MAX heterodimers by the competition of MXD, MNT, MGA proteins for MAX and these heterodimers for E-Box sequences at TSS. Moreover, MXD and MNT proteins can spread heterochromatin resulting in cell cycle arrest and differentiation [8,10,11]. MAX-like protein X (MLX), the cornerstone of a parallel network, can also heterodimerize with a subset of MXD proteins, *i.e.* MNT and MGA to repress metabolism and proliferation by binding to the Carbohydrate Response Element (ChoRE). MLX can also heterodimerize with MONDOA and CHREBP to activate transcription of these genes [10,11].

Deregulation of the *myc* paralogs leading to the accumulation of the corresponding gene products contribute to tumor growth by the sustained transcription of genes associated with growth, metabolism and proliferation [12–15]. On one hand, at such non-physiological levels, MYC is found to spill over or invade less specific sequences such as non-canonical E-Box and unrelated sequences at TSS of genes that contribute to the addiction of tumor cells [14,16]. On the other hand, the contribution of this invasion mechanism to tumorigenesis is debated because non-specific and weaker binding of the MYC/MAX heterodimer is not expected to lead to productive transcription [17].

MAX can also form a homodimeric b-HLH-LZ [18] that binds to chromatin and the E-Box sequences found at the promoters of MYC target genes [19,20]. When overexpressed *in cellulo*, MAX can act as an antagonist of the MYC/MAX heterodimer [21–23]. Moreover, an alternately spliced variant of MAX, delta-MAX (ΔMax), which is incapable of forming a homodimer, can still heterodimerize with MYC and bind DNA as a heterodimer is pro-oncogenic. In fact, and contrary to wildtype MAX, ΔMax activates the transcription of MYC target genes when over-expressed in cancer cells [24,25]. In addition, small molecules that can stabilize the MAX homodimer and its binding to chromatin have been shown to be efficacious to prevent the binding of the MYC/MAX heterodimer and its transcriptional activities in cancer cells overexpressing MYC [19,20]. Despite such evidence, the role of the MAX homodimer as an antagonist of MYC/MAX has been largely neglected in the literature perhaps because *max* was thought to be a housekeeping gene and hence not regulated.

The expression of MAX and MXD proteins is induced by TGF-b [26]. In accordance, whilst the expression of MYC is down-regulated by the positive TGF-b gradient observed along the crypt to villi axis (CVA) in the intestinal epithelium, MAX and MXD proteins expressions are upregulated [27,28]. Actually, while *c-myc* is maximally transcribed in the proliferative crypt cells, *max* and *mxd* genes are optimally expressed in the villi [27]. Hence, in the crypt cells, the level of MYC is larger than those of MXD and MAX. However, whereas MYC is barely detectable in the *villi*, MAX is more abundant than MYC and MXD proteins. Taken together, this suggests that the MYC/MAX heterodimer prevails in the crypt cells to promote proliferation, while the MAX/MAX homodimer followed by MXD/MAX heterodimers predominate in the *villi* to respectively halt proliferation and induce differentiation (Fig 1A).

**Figure 1.**
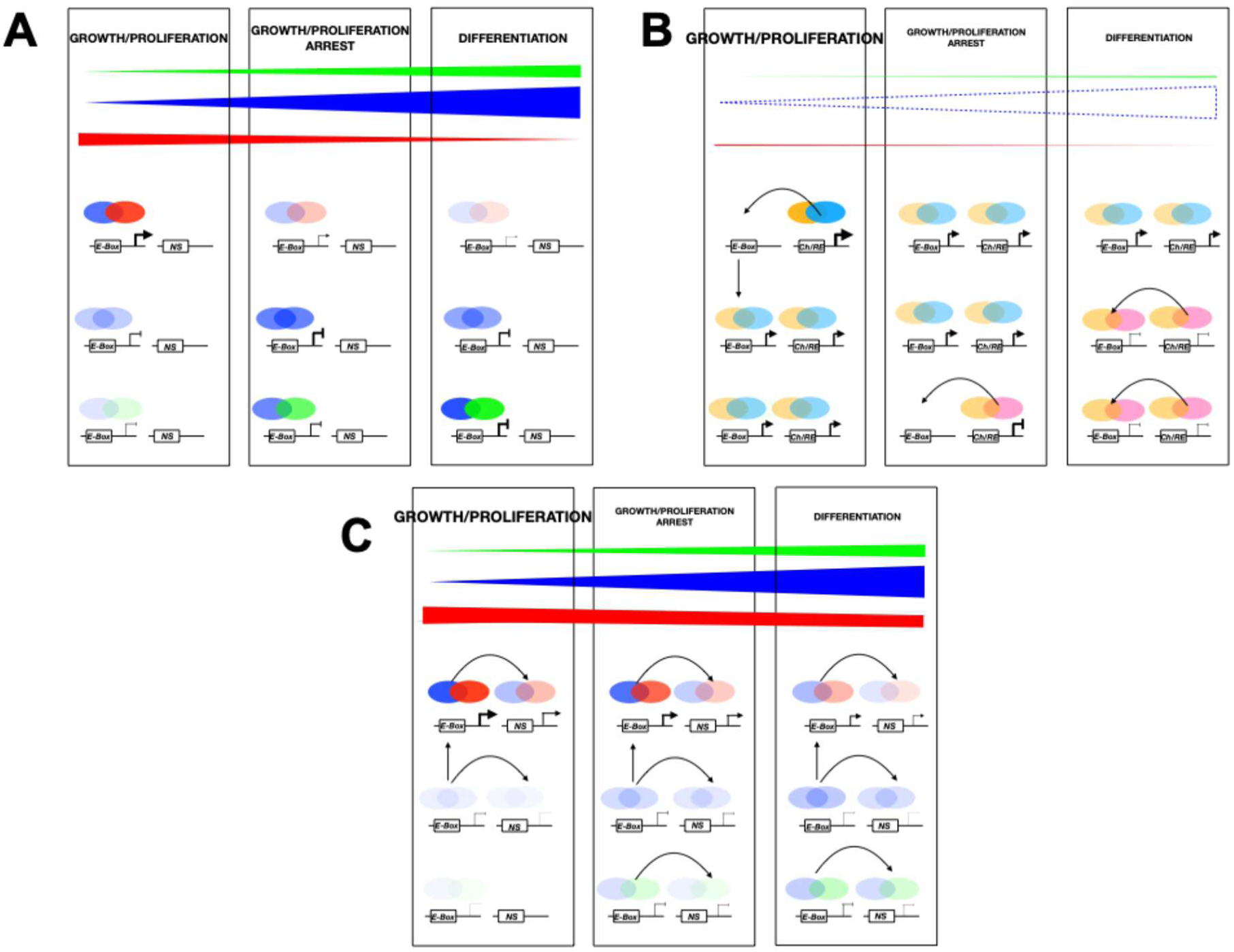
A. Regulation of the MYC/MAX/MAD network in the intestinal epithelium. MYC is in red, MAX is in blue and MAD is in green. Proteins expression levels are indicated by the height of the right-angle triangles of the corresponding colors. In accordance with the protein levels in the proliferating crypt cells, a predominant occupation of the E-Box by the MYC/MAX heterodimer occurs. The TGF-b dependent down-regulation of MYC and the up-regulation of MAX and MAD will lead to a shift from MYC/MAX heterodimers to the MAX/MAX homodimer onto the E-Box and to cell proliferation and growth arrests where differentiation begins, i.e. interface between the crypts and the villi [27]. Next, according to the observed protein levels the Mad/MAX heterodimer is expected to bind to the E-Box, spread heterochromatin and complete the differentiation of the villi. See text for further details. **B. Model for the effect of the ablation/inactivation of MAX and its tumor suppressor role**. In proliferating cells, the E-Box sequences that would be bound by MYC/MAX, MAX/MAX and MAD/MAX dimers are bound by MLX (orange) /MONDOA (light blue) and MLX/MNT (pink) [41]. In the absence of MAX, MYC and MXD proteins are rapidly degraded, hence the low levels of both families [41,57,60–62]. In this context, MLX/MONDOA activates MYC target genes to sustain proliferation. Although MLX/MNT binds to the E-Box, it does so to the expense of the Cho-RE. Hence, the extent of repression of both sets of genes is reduced and this is expected to delay cell proliferation and cell growth arrests as well as differentiation. **C. Model for the effect of variants of MAX with impaired affinity and/or specificity for the E-Box as homodimers.** These variants will preferentially partition as heterodimers onto DNA. This is expected to bring more MYC to the chromatin, protect it from degradation and enhance proliferation *vs.* a WT background. Variants that loose specificity are expected to bring MYC and MXD proteins onto non-specific sequences. Although non-specific binding of MYC to promoters may contribute to tumorigenesis, lack of binding of Mad proteins at the E-box is expected to delay cell proliferation and growth arrests as well as delaying differentiation.

Germline and somatic mutations in *max* leading to a truncated protein or complete lack of its expression have been reported in many cancers including those of neuro-endocrine origins such as SCLC, paraganglioma, including pheochromocytoma [29–39]. This raises the possibility that MAX may be a tumor suppressor. In an effort to gain mechanistic insights and validation of such a role for MAX, Augert *et al.* [40] and Freie *et al.* [41] have studied, *in cellulo* and *in vivo*, the impact of a MAX knockout in the early stages of SCLC and the truncation of the H2-LZ in the progression of neuroendocrine cancers, respectively. Results obtained by Augert *et al.* support the notion that the absence of MAX expression and the inability of MYC proteins, MXD proteins and MGA to heterodimerize and bind DNA, promotes, depending on the MYC and MXD proteins statuses, tumor growth. On the one hand, loss of MAX leads to the lack of binding of MYC to chromatin which leads to its degradation and the repression of its transcriptional programs sustaining growth and proliferation and, consequently, to growth and proliferation arrest. On the other hand, in cells where MXD/MAX heterodimers predominates over MYC/MAX heterodimers, the loss of MAX leads to the derepression of growth and metabolic genes and promote tumor growth [40]. Freie *et al.* [41] further demonstrated that *in vivo*, the ablation of the H2-LZ led to the expression of a truncated MAX protein incapable of homo- or heterodimerization and binding the E-Box. This caused the rapid degradation of MYC and allowed the binding of MYC target gene promoters by MLX/MONDOA and MLX/MNT heterodimers and to support metabolism and progression of neuro-endocrine tumors [41]. Hence the loss of expression of a functional MAX (capable of homodimerization, heterodimerization and DNA binding) leads to cell growth arrest when MYC levels are higher than MXD proteins and MGA or tumor progression when MYC protein levels are lower than MXD proteins and MGA.

These studies have shed unprecedented light into the effect of inactivating mutations that either abrogated expression of MAX or leads to the expression of a truncated and non-functional b-HLH-LZ in SCLC and neuro-endocrine cancers. However other SNPs (missense) leading to substitutions in the b-HLH-LZ are also described to be linked to similar phenotypes [29,30,34,35]. These mutations, although expected to modulate DNA binding of MAX as a homodimer and heterodimers, should not lead to complete inactivation. In this context, we believe it is important to address the impact of these mutations in order to have a broader view of all the possible ways that MAX could act as a tumor suppressor.

For example, amino acid substitutions in the basic region (*e.g.* H28R [42], E32K [43], R35C [30,42], R35P [44]) and in the HLH (*e.g.* R47W [42], R60W [45], R60Q [42,46–48]) have also been identified in a number of cancers including colon cancer, pheochromocytoma and in malignant neuroendocrine tumors [29,30,34,35,38,42–44]. Recently, the MAX^R60Q^ variant, a mutational hotspot, was found to occur *de novo* in three individuals and to lead to a complex disorder manifested by a macrocephaly, a polydactyly and delayed ophthalmic development [49]. The biophysical characterization of the MAX^R60Q^ b-HLH-LZ variant showed that it had a lesser affinity for the E-Box as a homodimer and a better affinity for the b-HLH-LZ of MYC that the WT b-HLH-LZ did. This mutation lead to preferential binding to the E-Box by the MYC/MAX^R60Q^ heterodimer compared to the wildtype. Accordingly, MYC target genes, including CyclinD2, were shown to be up-regulated when the MAX^R60Q^ variant was transfected in HEK 293 cells [49]. Accumulation of CyclinD2 caused by mutations in the mTOR pathway or directly in *CCND2* have already been linked to this developmental disorder [50–53].

Here, we continue on the characterization of three other variants found to occur in the basic region of MAX: E32K, R35C and R35P. All three homodimeric b-HLH-LZ (Max*) variants displayed a weaker affinity for the E-Box and are less effective than the wildtype at protecting against the binding of the heterodimeric MYC/MAX b-HLH-LZ. More precisely and similarly to the R60Q variant, Max*^R35P^ and Max*^R35C^ were found to have a higher affinity for the b-HLH-LZ of MYC (Myc*), a lower affinity for DNA and to lead, compared to the WT, to the preferential binding of the heterodimeric variants to the E-Box. Max*^E32K^ also bound DNA with a lesser affinity than the WT but does not have a better affinity for Myc*. Max*^E32K^ also lead to the preferential binding of the heterodimeric variant to DNA. However, whereas the Max*^R35C^ variant can still recognize the E-Box, Max*^E32K^ and Max*^R35P^ cannot discriminate between specific and non-specific DNA.

In the context where MYC predominates on MAX and on MXD, the MAX variants are expected to promote MYC binding to chromatin and stabilize it from degradation, thus sustaining the expression of MYC target genes. On the other hand, in the context of low MYC, and high MAX and MXD, as observed in the *villi* of the intestine, the sustained expression of these MAX variants will lead, compared to the non-pathogenic situation, to less homodimeric MAX and heterodimeric MXD/MAX at E-Box sequences. This is expected to delay cell growth arrest and differentiation and contribute to tumor growth. Altogether, our results offer additional mechanistic insights on the role of MAX as a tumor suppressor and further suggest that it is the MAX, in the homodimeric state, bound to the E-Box that acts as tumor suppressor.

## RESULTS

As described in **Fig. 2A and B**, R^35^ makes a non-specific salt bridge with the phosphodiester backbone. E^32^ and R^36^, which form specific H-Bonds are crucial for the mechanism of discrimination between an off-target sequence and the canonical CACGTG E-Box by the homodimeric and heterodimeric b-HLH-LZ of MAX [54,55]. Indeed, the carboxylate of E^32^ of one monomer accepts H-Bonds from the CA of one strand and the guanidino moiety of R^36^ to donate H-bonds to the G of the other strand. When in presence of a non-specific sequence, these interactions are not fulfilled, which destabilizes the helical state of the basic region. This leads to a complex with higher probability of leaving DNA to reengage in a new search process for the homodimeric b-HLH-LZ or even dissociate to heterodimerize with MYC and to find an E-Box and form a stable complex to activate transcription [17].

**Figure 2.**
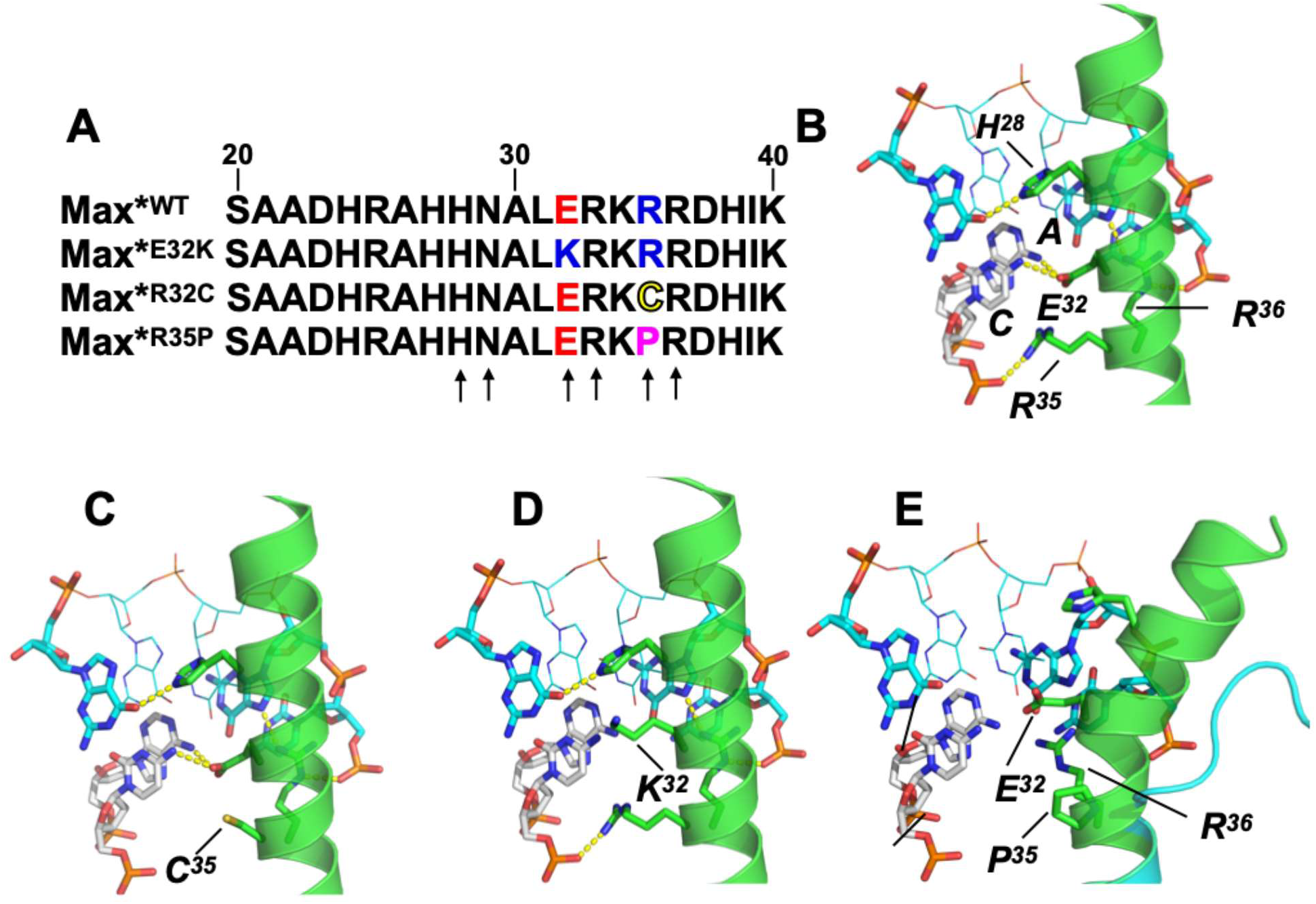
Structure schematics, specific and non-specific interactions dictating specificity and stability of binding of the basic region of MAX to the canonical (CACGTG) E-Box. **A.** Primary structure for the basic region of MAX and each of the variants. Positions making the most important contacts with the E-box are indicated by black arrows. Positions for the variants studied here are colored according to the Zappo colour scheme, following their physico-chemical properties: red for negative, blue for positive, magenta for proline and yellow for cysteine. **B.** The side chain (carboxylate) of E32 receives H-Bonds from the CA nucleobases in the leading strand (white carbon atoms). R35 and R36 make a salt bridges with phosphate groups while and the guanidino moiety of R36 makes a specific H-Bond with the nucleobase of the G in the strand of the reverse complement (cyan carbon atoms). **C.** The R35C mutation removes one non-specific salt-bridge at the interface of the complex. **D.** The aliphatic portion of the K side chain in the E32K variant is unable to accept the H-Bonds from the CA nucleobases and leads to the stabilisation of the complex and the helical structure of the basic region. **E.** In addition to removing a salt-bride, the Pro residue in the R35P kinks the path of the basic region, prevents the establishment of the specific H-Bonds mandatory for recognition of the E-Box and leads to unfolding of the helical state.

In **figure 3A**, we present the far-UV spectra recorded at 25°C of the Max*^E32K^, Max*^R35P^ and Max*^R35C^ along with that of Max*^WT^. As one can note, the spectra of Max*^R35P^ and Max*^R35C^ display more negative ellipticities than those Max*^WT^ and Max*^E32K^ indicating a slightly higher a-helical content and stability of the homodimeric states as judged by the [q]^MRW^ values at 222 nm ([q]_222_). [q]_222_ at 5°C ([q]_222_(5°C)) value for Max*^VL^, a stable mutant of the b-HLH-LZ of MAX with fully a folded HLH-LZ, but natively unfolded basic region in absence of DNA, was found to be around -22 000°·cm^2^·dMol^-1^ [3]. The fact that the [q]_222_(5°C) values are less negative suggests that all four constructs form less helical dimers and/or exist as an equilibrium between a population of dimeric state (P_D_) and a population of unfolded monomer (P_M_). Simulation of the denaturation curves to a two-state denaturation of a native dimeric state into two unfolded monomers [3] indicates P_M_(5°C) of 0.15 for Max*^WT^, 0.12 for Max*^E32K^, 0.07 for Max*^R35C^ and 0.06 Max*^R35P^ with [Q_N_]_222_(0°C) value of -20 000°·cm^2^·dMol^-1^ for the native homodimers (See supplementary material). This values also suggest that the homodimers are not as helical as Max*^VL^ which has a stabilized and fully helical leucine-zipper. Congruently, the homodimeric LZ of MAX is not fully helical at 0°C [56]. In accordance with their lower the P_M_(5°C), the temperature denaturation curves of Max*^R35P^ and Max*^R35C^ are slightly shifted towards higher T° values indicating a slightly higher stability for these homodimers (**see Table S2**). As documented elsewhere [3], the stability of the HLH of Max*^WT^ is limited by quaternary electrostatic repulsions between R and K residues at the interface of the b-HLH, one of these being R^35^. Hence, replacing R^35^ by a P or a C will reduce these repulsions and might account for the slight increase in apparent stability of these homodimers.

**Figure 3.**
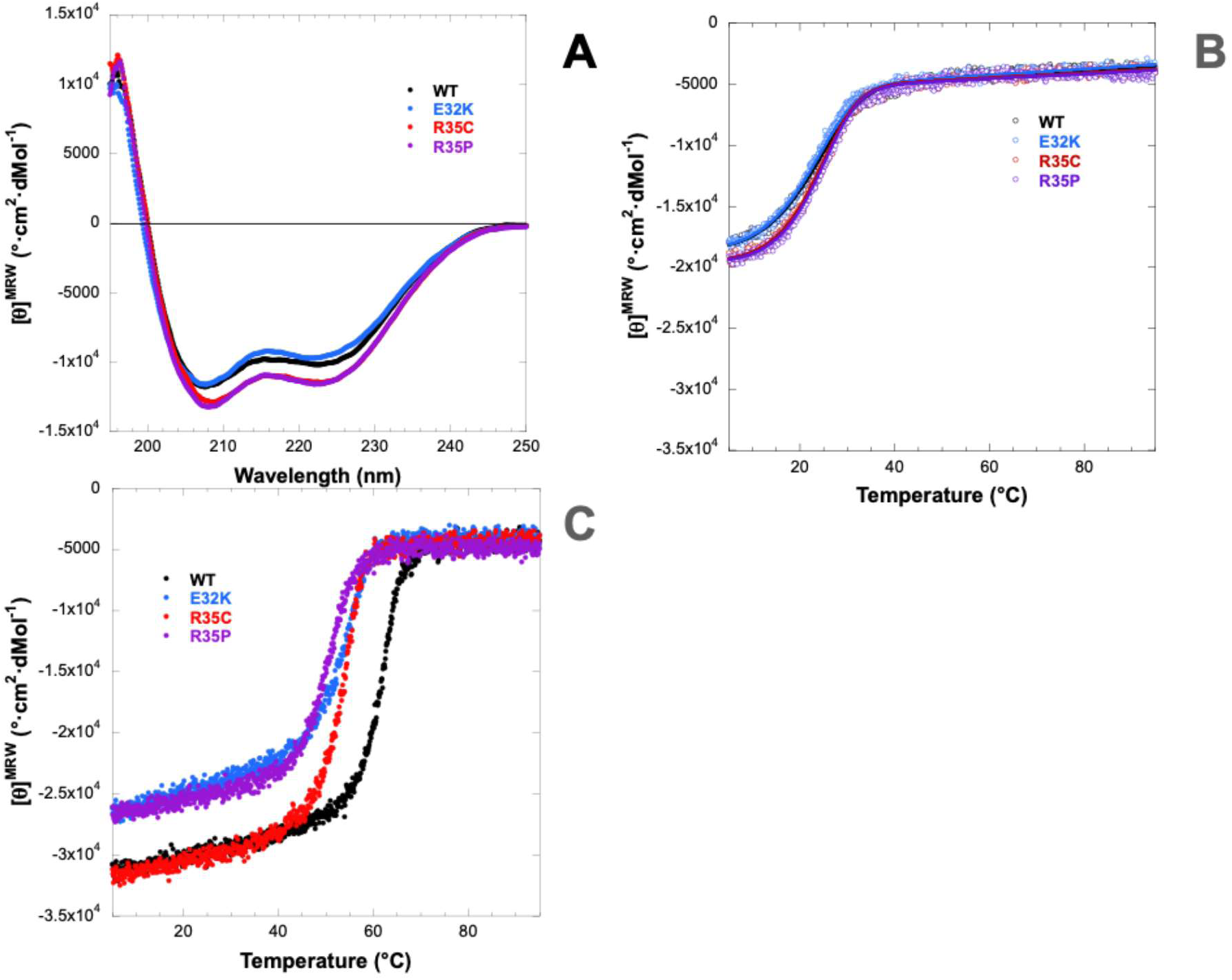
**A.** Far-UV CD of Max*^WT^ (black), Max*^E32K^ (blue), Max*^R35P^ (purple) and Max*^R35C^ (red) recorded at 20 mM (monomer units), 25°C and pH 6.8. **B.** Corresponding temperature denaturation recorded under the same conditions. Same color coding as in **A**. **C.** Temperature denaturation Max*^WT^, Max*^E32K^, Max*^R35P^ and Max*^R35C^ recorded at 20 mM (monomer units) in the presence of 20 mM (duplex units) of a canonical E-Box probe at 25°C and pH 6.8. The DNA signal was subtracted. Same color coding as in **A**. The solid lines represent the simulation of a two-state denaturation between a native homodimer and two unfolded monomers. The parameters of the simulations are presented in Table S1.

We next sought to evaluate the relative stability of the corresponding homodimeric complexes with the canonical E-Box (**Fig. 3C**). As expected, both the ellipticity and the apparent T° of the Max*^WT^ are optimal in the presence of the canonical E-Box. As described above, in the presence of the E-Box, the side chain of E^32^ and R^36^ can accept and donate specific H-Bonds (**Fig. 2B**) and this allows for the complete folding of the basic region into a a-helix and the formation of a specific and optimally stable complex. One can also see that Max*^R35C^, which lacks one non-specific salt-bridge (**Fig. 2C**), can achieve the same ellipticity and hence form the specific H-Bonds but forms a less stable complex with the E-Box. However, the apparent helical content and T° obtained for Max*^E32K^ are both lower. Compared to the E, side chain of K has a larger hydrophobic surface and a positively charged amino group (**Fig. 2D**). Hence Max*^E32K^ cannot accept the H-Bonds from the CA and allow for the stabilization of a fully helical complex. Likewise, and despite its native basic region, the Max*^R35P^ construct has the lowest helical content and apparent stability in the presence of the E-Box. As shown in **Fig. 2E**, the presence of a P at position 35 kinks the helical basic region away from the Hoogsteen edge and prevents the establishment of specific (and non-specific) contacts between E^32^ and the CA pairs in the E-Box sequence. As a result, the basic region is most likely unfolded in both complexes hence explaining the lowest stability amongst studied variants.

In **Figure 4**, we explore the capacity of the Max* constructs to discriminate between the canonical E-Box and a scrambled version (nsDNA) which does not allow for the formation of specific H-Bonds by E^32^ and R^36^. We note the expected decrease in helical content and stability when the Max*^WT^ is in presence of the nsDNA compared to the E-Box (**Fig. 4A**). As a proxy for discrimination, we estimated the differences in the apparent T° between the E-Box and nsDNA bound constructs. Most interestingly, whereas the ΔT° for the Max*^WT^ construct is 7°C, no apparent ΔT and difference in helical content are observed for the Max*^R35P^ construct (**Fig. 4B**). This confirms that the Max*^R35P^ construct is unable to discriminate between specific and non-specific DNA. This can be explained by the kink induced by the P which prevents the basic region to bury itself into the major groove and probe for specific interactions as described elsewhere [55]. One can also note that the lack of non-specific electrostatic interaction between R35 and the phosphodiester backbone diminishes discrimination of the Max*^R35C^ construct with a ΔT° of 3.5° (**Fig. 4C**). Finally, we also observe a lack of discrimination for the Max*^E32K^ (**Fig. 4D**). In this case, the binding to the scrambled E-Box seems to be slightly favored (ΔT°=-0.5). It has to be noted that one finds a TC in the scrambled E-Box instead of a CA of the leading strand of the E-Box. This leads to the possibility of formation of hydrophobic interaction between the methyl group on the T nucleobase and the hydrophobic portion of the K32 side chain and may explain the slight preference observed (not shown).

**Figure 4.**
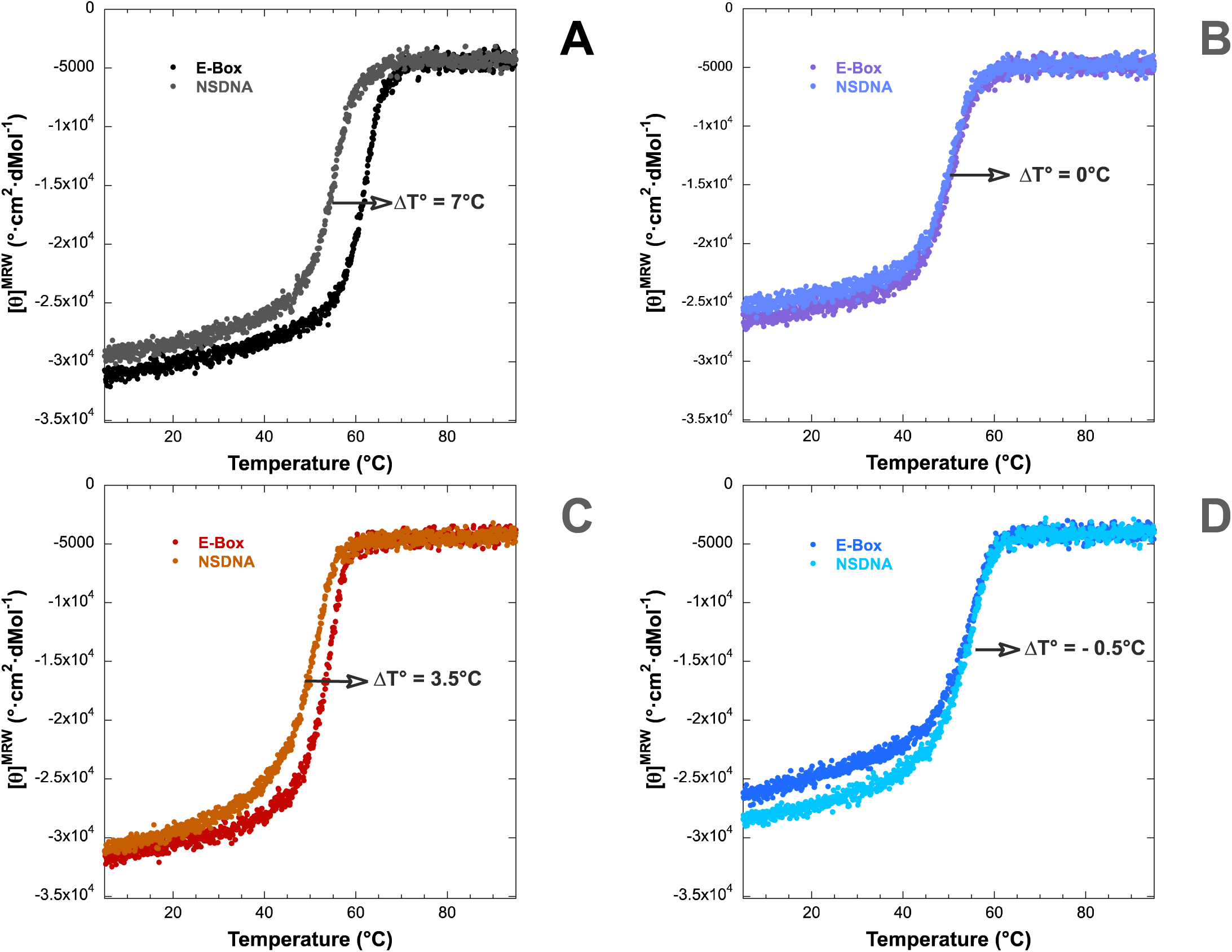
Discrimination between the canonical E-Box and the non-specific DNA probe. **A.** Temperature denaturation of Max*^WT^ (20 mM; monomer units) in the presence of the E-Box and the non-specific probes (20 mM; duplex units). **B.** Temperature denaturation of Max*^WT^ (20 mM; monomer units) in the presence of the E-Box and the non-specific probes (20 mM; duplex units). C. Temperature denaturation of Max*^WT^ (20 mM; monomer units) in the presence of the E-Box and the non-specific probes (20 mM; duplex units). **D.** Temperature denaturation of Max*^WT^ (20 mM; monomer units) in the presence of the E-Box and the non-specific probes (20 mM; duplex units). The DNA signal was subtracted.

We next moved on to evaluate the effect of these mutations on the heterodimerization with Myc*. We present in **Figure 5**, the temperature denaturation of Myc* (at 10µM) in the presence of Max*^E32K^ (blue), Max*^R35P^ (purple) and Max*^R35C^. As described above, the WT and Myc*/Max^E32K^ reach the expected [q]_222_(5°C) value for a fully and natively folded heterodimeric b-HLH-LZ. However, the Myc*/Max*^R35P^ and Myc*/Max*^R35C^ possess a lower and higher [q]_222_(5°C) value, respectively. On the one hand, the lower value of Myc*/Max*^R35P^ could be explained by the fact that the destabilization of the beginning of Helix 1 by the P may have a long-range effect. Indeed, this region is involved in quaternary interaction with the beginning of Helix 2 of MYC. Hence it is likely that the presence of a P, instead of a R, may reduce the helical content of the Myc*/Max*^R35P^ by weakening the secondary structure of the MAX H1 and quaternary interactions. On the other hand, the apparent helical content of the Myc*/Max*^R35C^ is the largest. In contrary to P, the presence of a C will not reduce the helical content of the H1 and interaction with H2 of Myc* and coupled to the reduction of quaternary repulsions it is somewhat expected that this heterodimer is also more stable than the WT. In order to quantify and validate this, the denaturation curves were simulated with a two-state model for denaturation of a heterodimer into two unfolded monomers. The consistent results of the simulation can be found in **Table S3**.

**Figure 5.**
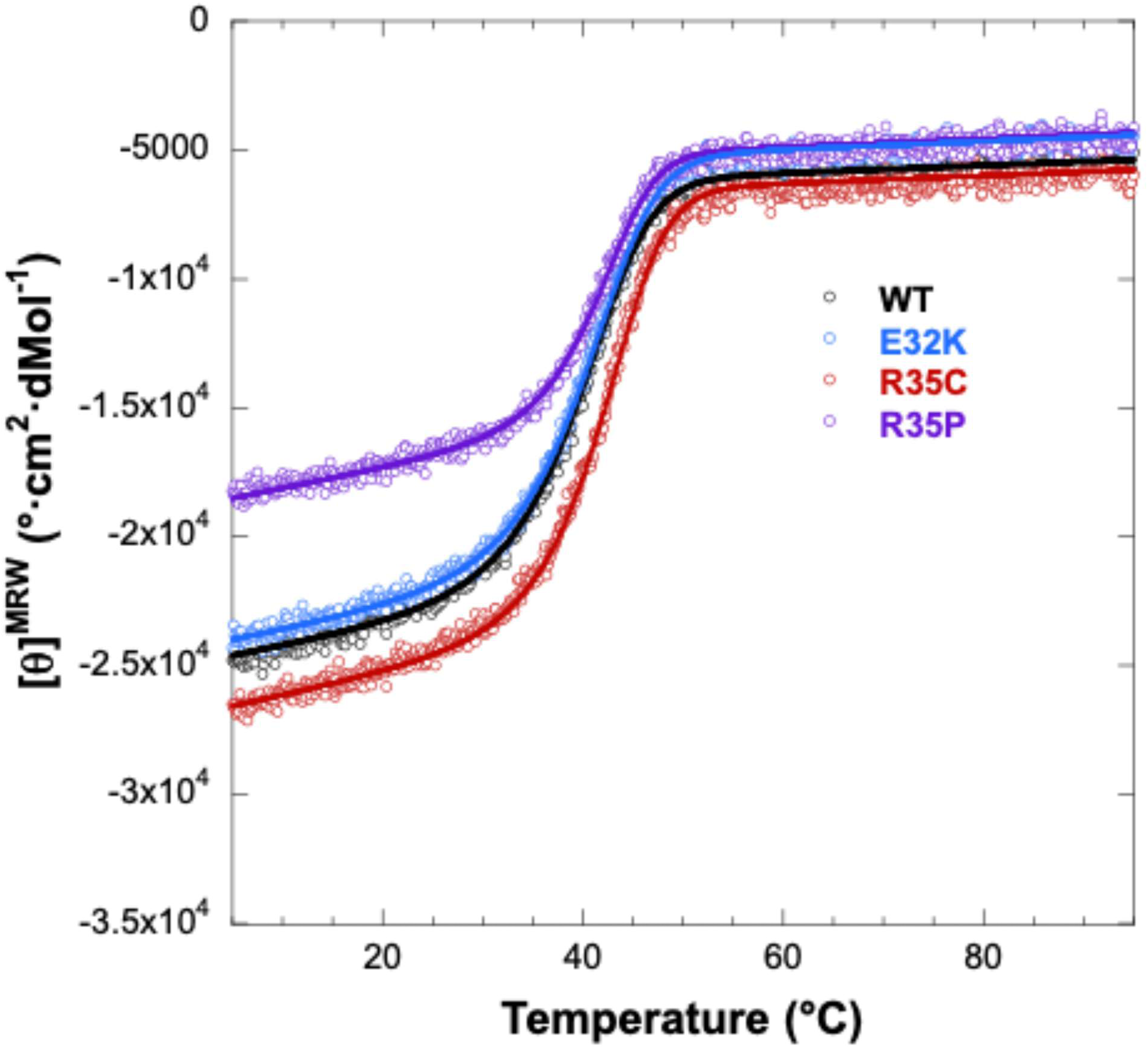
Heterodimerization between Max*^WT^ and its variants with Myc*. Temperature denaturation of Myc* (10 mM, monomer units) in the presence of Max*^WT^ (10 mM, monomer units: black), Max*^E32K^ (10 mM, monomer units: blue), of Max*^R35P^ (10 mM, monomer units: purple) and Max*^R35C^ (10 mM, monomer units: red). The solid lines represent the simulation of a two-state denaturation between a native heterodimer and two unfolded monomers. The parameters of the simulations are presented in Table S2.

These results are very important with respect to the role of MAX in controlling the binding of MYC at E-box sequences. Indeed, the MAX homodimer bound at E-Box sequence competes with the binding of the heterodimer [19,20]. Hence loss of affinity of the MAX homodimer for the E- box sequence and an increase in affinity for the b-HLH-LZ of MYC should favor a higher occupancy of the E-Box by the MYC/MAX heterodimer for a fixed level of MYC. In order to investigate such a phenomenon, we have studied the relative proportions of the Max* constructs in homodimeric and heterodimeric complexes with the E-Box. In order to do so we incubated the different constructs at 15 mM in the presence of 10 mM Myc* and 10 mM of the double-stranded E-Box and proceeded to record temperature induced denaturations. This ratio is a realistic representation of the relative levels of MYC and MAX in the crypt cells of the intestinal epithelium (**Fig. 1A**). In this approach the amount of relative dimeric populations bound to the E-Box can be estimated by the relative amplitudes of their denaturation appearing with a 5000 nM excess of the Max* constructs as described by the arrows in **Fig. 6**.

**Figure 6.**
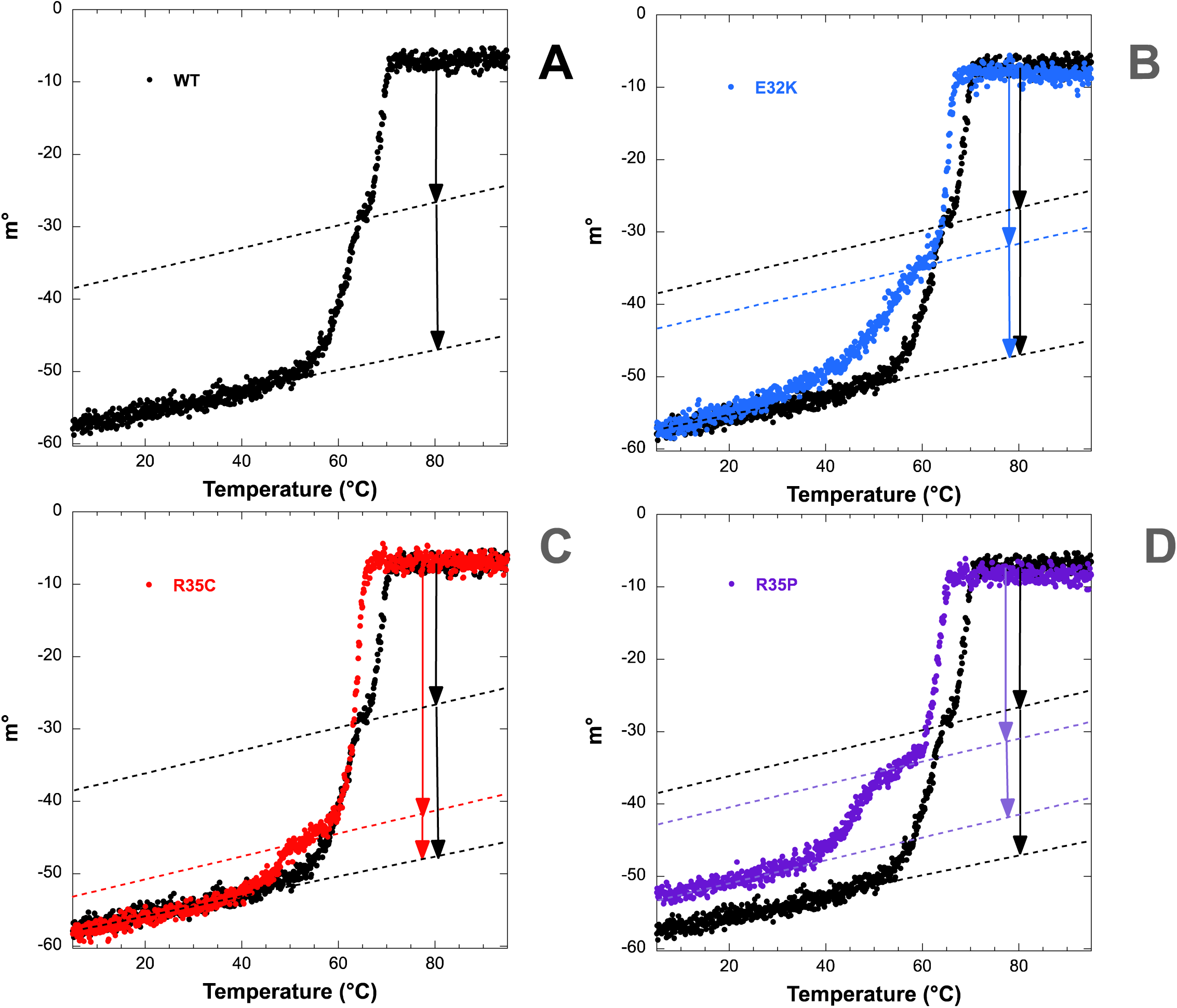
Max*^WT^ construct is the most efficient construct for preventing the binding of the heterodimeric Myc* to the E-Box. **A.** Temperature denaturation of Myc* (10 mM, monomer units) and Max*^WT^ (10 mM, monomer units) in the presence of the E-Box probe (10 mM duplex units). Note the residual presence of the homodimeric Max*^WT^:E-Box complex. **B.** Temperature denaturation of Myc* (10 mM, monomer units) and Max*^E32K^ (10 mM, monomer units in blue) in the presence of the E-Box probe (10 mM duplex units). **C.** Temperature denaturation of Myc* (10 mM, monomer units) and Max*^R35C^ (10 mM, monomer units in red) in the presence of the E-Box probe (10 mM duplex units). **D.** Temperature denaturation of Myc* (10 mM, monomer units) and Max*^R35P^ (10 mM, monomer units in purple) in the presence of the E-Box probe (10 mM duplex units). The DNA signal was subtracted.

Figure 6A displays the relative populations of the Max*^WT^/Max*^WT^ (arrow at lower apparent T°) and Myc*/Max*^WT^ (arrow for the transition at higher T°). Those of Max*^E32K^, Max*^R35C^ and Max*^R35P^ are presented on **Figures 5B, C and D**, respectively. As expected, the Max*^WT^ homodimer is the most abundant dimer bound to the E-Box and as such leads to the lowest population of all the Myc*/Max* dimer on the cognate DNA sequence. In other words, because Max*^WT^ is more abundant and has the highest affinity (*vs*. variants) for the E-Box as a homodimer it is the best at preventing the E-Box from being bound by Myc* as a heterodimer. Conversely, owing to their lower affinity for the E-Box as homodimers, Max*^E32K^ (**Fig. 5B**), Max*^R35P^ (**Fig. 5C**) and Max*^R35C^ (**Fig. 5D**) preferentially partition on the E-Box sequence as heterodimers. Note that the Max*^R35C^ construct is the least proficient homodimer for blocking MYC from binding to the E-Box. This is likely due to the fact that in addition to having a lower affinity for the E-Box compared to the WT as a homodimer, it is the construct with the highest affinity for the b-HLH-LZ of MYC.

## DISCUSSION

When upregulated MYC proteins (c-, N- and L-) are prevailing oncoproteins sustaining all aspects of tumorigenesis and tumor growth. Inactivating and missense mutations in MAX, their obligate partner, are linked many cancers and other diseases. This has led to the concept that MAX could be a tumor suppressor. In support, the inactivation of Max though truncation of its H2-LZ was shown to lead to the onset of neuro-endocrine tumors in mice [40,57]. This truncated Max is unable to bind to the E-Box as a homodimer and prevented the binding of MYC to chromatin and consequently led to its proteasomal degradation. In the absence or low levels of MYC, tumor growth was revealed to be sustained by the binding of MLX/MONDOA and MLX/MNT to vacated E-Box sequences. In such a scenario, MLX/MONDOA is able to rescue the transcription of MYC target genes. However, whereas MLX/MNT binding at E-Box sequences could lead to some repression of MYC target genes this is expected to be done at the expense of the derepression of metabolic genes driven by the Cho/RE and to promote proliferation, growth and delay differentiation (**Fig. 1B**). [10,40,57]

This model offers a mechanism to rationalize why the inactivation of MAX can lead to tumorigenesis. However, variants leading to missense mutations are also associated with cancers such as pheochromocytomas, paragangliomas and other lesions as well as an overgrowth syndrome [29,30,34,35,38,42–48]. Such mutations are expected to lead to the expression of variants with altered dimerization and DNA binding behaviour and not complete inactivation. Recently the most recurrent variant of MAX found in the Loop region, MAX^R60Q^, associated with gliomas, carcinomas and other lesions [42,45,48,58], was found to occur *de novo* in newborns and to cause overgrowth phenotypes such as macrocephaly and polydactyly [49](Harris et al. 2024). It was demonstrated that the MAX^R60Q^ b-HLH-LZ (Max*^R60Q^), has a lower affinity as a homodimer for the E-Box sequence that the WT and a higher affinity for the b-HLH-LZ of MYC (Myc*). This was shown, *in vitro*, to result in a more sustained occupation of the E-Box by the Max*^R60Q^/Myc* heterodimer compared to the WT. Accordingly, *in cellulo*, MYC target genes such as Cyclin D2 were shown to be overexpressed, compared to the WT, when Max^R60Q^ was transiently transfected in HEK 293 cells. In this scenario, it is reasonable to assume that in combination with MYC deregulation the Max^R60Q^ and the other variants with missense mutations may play a pro-oncogenic role in cancer malignancies. Here, we extended the characterization of such MAX variants, *i.e.* Max*^E32K^, Max*^R35P^ and ^R35C^. These are all located in the basic region and positions 32 and 35 which are involved in the specific and non-specific interactions with the E-Box, respectively (**Fig. 2**). Our results demonstrate that each variant also favors the formation of the heterodimeric b-HLH-LZ complex on the E-Box sequence. This is promoted by the loss affinity for DNA as homodimers by all three variants and an increase in affinity for the b-HLH-LZ of MYC for Max*^R35P^ and Max*^R35C^ specifically. As shown in **Fig. 1C**, this favorable binding of the heterodimers to the E-Box coupled to the increased availability is expected to stabilize MYC, promote proliferation and growth and delay differentiation.

Although the Max*^R35C^ homodimer can still discriminate between the E-Box and a scrambled E-Box, the Max*^E32K^ and Max*^R35P^ cannot. For the latter, we show that this is caused by the P at position 35 kinking the basic region away for the major groove and preventing it from recognizing the E-Box (**Fig. 2E**). For the former, the K side chain cannot receive the key H-Bonds from the CA in the E-Box (**Fig. 2B and D**). Congruently, replacement of the conserved residues involved in the recognition of the E-Box in the basic region of MYC (*i.e.* H359, E363) profoundly impacted the recognition of the E-Box and the genome-wide localisation of MYC [17]. In fact, this mutant of MYC was shown to still be able to bind the E-Box sequences but also half-E-Box sequences and to other unrelated sequences. This underscores the importance of the conserved E and also suggests that the recognition of the E-Box and the half-E-Box rely on the basic region of MAX. While the invasion of upregulated MYC to such non-specific sites has been suggested to play an amplifier role that contributes to sustain oncogenic transcriptional programs [14,15,59], this notion is being debated. It is argued that such non-specific leads to DNA bound MYC/MAX heterodimers may not be stable enough to reside on chromatin long enough to sustain active transcription. This actually suggests that the MAX variants/MXD heterodimers will also bind to non-related sequences and in turn become less efficient at repressing the expression of MYC target genes. As depicted in **Fig. 1C**, this is expected to further delay differentiation and contribute to tumor growth.

In conclusion, cancer malignancies have been associated with inactivating and missense mutations in MAX. This has led to the notion that MAX has a tumor suppressor function. In support of this notion, inactivation of MAX has been shown to promote neuro-endocrine tumors in mice by leading to the degradation of MYC and the replacement of the MYC/MAX heterodimer on chromatin by heterodimeric b-HLH-LZ transcription factors from the MLX network, which are able to sustain cell growth and delay differentiation of neuro-endocrine tissues. Here, we show that *in vitro*, potentially oncogenic variants with missense mutation in the basic region of the b-HLH-LZ of MAX (E32K, R35C and R35P) lose affinity for DNA as homodimers and lead to the preferential binding of the MYC/MAX heterodimeric b-HLH-LZ to the E-Box. Hence, these variants are expected to be prooncogenic by stabilizing oncogenic levels of MYC on chromatin.

As discussed, our results support the concept that the homodimeric state of MAX bound to DNA can play a tumor suppressor role by limiting the levels of MYC proteins and members and heterodimers from the MLX network bound to the E-Box [20].

