## Supplementary information for "Basic region variants of the MAX b-HLH-LZ preferentially form heterodimers with the MYC b-HLH-LZ to bind the E-box rather than binding as homodimers"

**Table S1: Primers used to clone the Max variants**

| Variant | Primers | |
|---|---|---|
| E32K | Forward | 5'-GACAAACGGGCTCATCATAATGCACTGAAACGAAAACGTAAGGACCAC-3' |
| | Reverse | 5'-CAAACTGTGAAAGCTGTCTTTGATGTGGTCCCTACGTTTCAGTGCAATATGATG-3' |
| R35C | Forward | 5'-GGGCTCATCATAATGCACTGGAACGAAAATGTAGGGACCACATCAAAGAC-3' |
| | Reverse | 5'-CAAACTGTGAAAGCTGTCTTTGATGTGGTCCCTACATTTTCGTTCCAGTGC-3' |
| R35P | Forward | 5'-GGGCTCATCATGCACTGGAACGAAAACCTAGGGACCACATCAAAGAC-3' |
| | Reverse | 5'-CAAACTGTGAAAGCTGTCTTTGATGTGGTCCCTAGGTTTTCGTTCCAGTGC-3' |

**Table S2 : Thermodymical parameters of the simulation of the temperature denaturations of the WT and variants of the b-HLH-LZ of Max at 20 μM with a two-state transition model of the dissociation/unfolding between a homodimer and two unfolded monomers**

| | $\Delta H°_u$ (T°)[1] | T° (°C) | $K_D$ (5°C) | $P_M$ (5°C) | $K_D$(25°C) | $P_M$(25°C) |
|---|---|---|---|---|---|---|
| WT | 40.0 | 23.0 | $1.0\cdot10^{-6}$ | 0.15 | $5.5\cdot10^{-5}$ | 0.67 |
| $E^{32}K$ | 44.0 | 23.0 | $6.1\cdot10^{-7}$ | 0.12 | $5.7\cdot10^{-5}$ | 0.68 |
| $R^{35}C$ | 53.0 | 24.5 | $1.8\cdot10^{-7}$ | 0.07 | $4.1\cdot10^{-5}$ | 0.62 |
| $R^{35}P$ | 54.0 | 24.7 | $1.5\cdot10^{-7}$ | 0.06 | $3.7\cdot10^{-5}$ | 0.61 |

[1]Values in $kcal\cdot mol^{-1}$

**Table S3 : Thermodymical parameters of the simulation of the temperature denaturations of the WT c-Myc* at 10 μM with 10 μM of the WT and variants of the b-HLH-LZ of Max with a two-state transition model of the dissociation/unfolding between a heterodimer and two unfolded monomers**

| | $\Delta H°_u$(T°)[1] | T° | $K_D$(5°C) | $P_M$(5°C) | $K_D$(25°C) | $P_M$(25°C) |
|---|---|---|---|---|---|---|
| WT | 80 | 41.5 | $6.3\cdot10^{-11}$ | $1.3\cdot10^{-3}$ | $5.7\cdot10^{-8}$ | $3.7\cdot10^{-2}$ |
| $E^{32}K$ | 80 | 41.7 | $5.7\cdot10^{-11}$ | $1.3\cdot10^{-3}$ | $4.8\cdot10^{-8}$ | $3.4\cdot10^{-2}$ |
| $R^{35}C$ | 90 | 43.5 | $4.6\cdot10^{-12}$ | $3.4\cdot10^{-4}$ | $1.3\cdot10^{-8}$ | $1.7\cdot10^{-2}$ |
| $R^{35}P$ | 99 | 43.0 | $7.4\cdot10^{-13}$ | $1.4\cdot10^{-4}$ | $5.7\cdot10^{-9}$ | $1.2\cdot10^{-2}$ |

[1]Values in $kcal\cdot mol^{-1}$

The baseline for the native and dimeric (N) as well as unfolded and monomeric states (U) were considered linear and given by the following functions:

$$[\theta_N]_{222}(T) = [\theta_N]_{222}(0°C) + m(\theta_{222}/°C) \cdot T(°C)$$

$$[\theta_U]_{222}(T) = [\theta_U]_{222}(0°C) + m_U(\theta_{222}/°C) \cdot T(°C)$$

Where $[\theta_N]_{222}(T)$ and $[\theta_U]_{222}(T)$ are the linear temperature dependent mean residue molar ellipticity of the N and U states, respectively. $[\theta_N]_{222}(0°C)$ and $[\theta_U]_{222}(0°C)$ are the corresponding abscissas at the origin and $m_N$ and $m_U$ the slopes of $[\theta_N]_{222}(T)$ and $[\theta_U]_{222}(T)$.

The values used for the simulations of the temperature denaturation curves of the homodimers were:

| | WT | $E^{32}K$ | $R^{35}C$ | $R^{35}P$ |
|---|---|---|---|---|
| $[\theta_N]_{222}(0°C)$ | -20 000 | -19 600 | -20 000 | -20 000 |
| $[\theta_U]_{222}(0°C)$ | -5 600 | -5 750 | -5 700 | -5 750 |
| $m_N$ | -75 | -50 | -50 | -50 |
| $m_U$ | 22 | 25 | 20 | 20 |

And those for the simulations of the temperature denaturation curves of the heterodimers were:

| | WT | $E^{32}K$ | $R^{35}C$ | $R^{35}P$ |
|---|---|---|---|---|
| $[\theta_N]_{222}(0°C)$ | -27 000 | -24 700 | -27 000 | -18 900 |
| $[\theta_U]_{222}(0°C)$ | -6 800 | -6 600 | -7 200 | -5 850 |
| $m_N$ | 75 | 90 | 85 | 78 |
| $m_U$ | 15 | 20 | 15 | 16 |

As described elsewhere (JFN 2023), the actual curves $[\theta]_{222}(T)$ were simulate according to:

$$[\theta]_{222}(T) = (1-P_U(T)) \cdot [\theta_N]_{222}(T) + P_U(T)) \cdot [\theta_U]_{222}(T)$$

Where $P_U$, the population of the U state is given by:

$$P_U(T)=K_U(T)/(1+K_U(T))$$

and $K_U(T)$, the temperature dependent equilibrium by:

$$K_U(T) = e^{(-\Delta G^\circ_U(T)/RT)}$$

$\Delta G^\circ_U(T)$ is the temperature dependent change in the standard Gibbs free energy between the U and N states, R the gas constant in kcal and T the absolute temperature in K.

Finally,

$$\Delta G^\circ_U(T) = (\Delta H^\circ_U(T^\circ) + \Delta C^\circ_{p,U} \cdot (T-T^\circ))$$
$$- T \cdot ((\Delta H^\circ_U(T^\circ)/T^\circ + \Delta C^\circ_{p,U} \cdot \ln (T/T^\circ)) + (R\cdot \ln(1.66\cdot pT)))$$

Where $\Delta H^\circ_U$ is the change in standard molar enthalpy between the U and N states and pT is the total concentration of monomers in M. A change standard heat capacity at constant pressure between the U and N states ($\Delta C^\circ_{p,U)}$ of 0.9 kcal·mol$^{-1}$·K$^{-1}$ was used is all simulation.
